# Tissue-specific transcriptome for *Poeciliopsis prolifica* reveals evidence for genetic adaptation related to the evolution of a placental fish

**DOI:** 10.1101/289488

**Authors:** Nathaniel K. Jue, Robert J. Foley, David N. Reznick, Rachel J. O’Neill, Michael J. O’Neill

**Affiliations:** Institute for Systems Genomics and Department of Molecular and Cell Biology, University of Connecticut, Storrs, CT 06269; School of Natural Sciences, California State University, Monterey Bay, Seaside, CA 93933; Department of Biology, University of California, Riverside, CA 92521

**Author notes:** Corresponding author: Institute for Systems Genomics and Department of Molecular and Cell Biology, University of Connecticut, Storrs, CT 06269, USA. Data deposition: All read data was deposited in the NCBI SRA database under the following accession numbers: SRR1639275, SRR1640127, SRR1640137, SRR1640160, SRR1640171, SRR1640200, SRR1640209, SRR1640216, and SRR1640219 under the BioProject PRJNA266248. This Transcriptome Shotgun Assembly project has been deposited at DDBJ/EMBL/GenBank under the accession GBYX00000000.1 and the same BioProject.

**Keywords:** Transcriptome, Positive Selection, Gene Expression, Placenta, Fish

## Abstract

The evolution of the placenta is an excellent model to examine the evolutionary processes underlying adaptive complexity due to the recent, independent derivation of placentation in divergent animal lineages. In fishes, the family Poeciliidae offers the opportunity to study placental evolution with respect to variation in degree of post-fertilization maternal provisioning among closely related sister species. In this study, we present a detailed examination of a new reference transcriptome sequence for the live-bearing, matrotrophic fish, *Poeciliopsis prolifica*, from multiple-tissue RNA-seq data. We describe the genetic components active in liver, brain, late-stage embryo, and the maternal placental/ovarian complex, as well as associated patterns of positive selection in a suite of orthologous genes found in fishes. Results indicate the expression of many signaling transcripts, “non-coding” sequences and repetitive elements in the maternal placental/ovarian complex. Moreover, patterns of positive selection in protein sequence evolution were found associated with live-bearing fishes, generally, and the placental *P. prolifica*, specifically, that appear independent of the general live-bearer lifestyle. Much of the observed patterns of gene expression and positive selection are congruent with the evolution of placentation in fish functionally converging with mammalian placental evolution and with the patterns of rapid evolution facilitated by the teleost-specific whole genome duplication event.

## INTRODUCTION

The study of the placenta provides insight into the evolutionary relationships of biological phenomena such as complexity, live-birth and genetic conflict. A great deal of research has focused on the function and development of mammalian placentas, uncovering the unique regulatory, genetic, and evolutionary nature of this structure. Studies of gene regulation in the mammalian placenta show a suite of unique features including genomic imprinting (Bressan *et al*. 2009), non-coding RNAs (Koerner *et al*. 2009), and DNA methylation and histone-modification mediated transcription (Maltepe *et al*. 2010). The placenta also has been shown to be a tissue that utilizes genes derived from the co-option of retroelements for unique functional purposes (Lavialle *et al*. 2013). Additionally, the placenta has been used as a model for examining the evolution of tissue-specific novelties, such as newly derived cell-types (Lynch *et al*. 2011), placental variation among eutherian mammals (Carter and Mess 2007), and genomic imprinting related to viviparity (Renfree *et al*. 2013).

Placentation is typically studied in mammals, but fish present a compelling study system for examining contributing factors to the evolution of this complex organ. The Neotropical fish family Poeciliidae is comprised of approximately 200 species, all of which, with one exception, give live birth. The majority of these poeciliids are lecithotrophic (i.e. yolk-feeding), wherein eggs provide all necessary nutrients to support the embryo through development to birth.

However, placenta-like structures that permit post-fertilization maternal provisioning have evolved independently in multiple poeciliid lineages, specifically within certain groups such as species in the genus *Poeciliopsis*, within the last 750,000 years (Reznick *et al*. 2002). Unlike comparisons between eutherian and marsupial mammals, who last shared an ancestor with their non-placental monotreme counterparts (i.e. the egg-laying platypus and echidnas) ∼200 million years ago (Meredith *et al*. 2011), species within *Poeciliopsis* offer the opportunity to investigate more “recent” changes leading to viviparity and placentation. The relatively recent adaptation of placentation has resulted in wide variation among *Poeciliopsis* species with respect to the extent of maternal provisioning. The extent of maternal investment across species ranges from highly matrotrophic (i.e. placentotrophic) to lecithotrophic, including intermediate, or “partial”, placental species. These transitional states and independent evolutionary events make this system particularly powerful for examining factors contributing to the evolution of placentation (see Pollux *et al*. 2009 for review).

Although fish placentas exhibit functional convergence, they are diverse in structure, with poeciliid placentas bearing features distinct from mammalian placentas. In poeciliids, the maternal portion of the placenta is derived from the ovarian follicle. Fertilization occurs within the ovarian follicle wherein the embryo will subsequently develop. Within placental *Poeciliopsis* species, nutrient exchange occurs across an enlarged pericardial sac that contributes to a large, highly vascularized belly sac (Turner 1940). In the closely related poeciliid species *Heterandria formosa*, functionally convergent placental structures are notably divergent in structure; the aforementioned sac structure covers regions more anterior on the developing embryo (Turner 1940). While specializations to the follicular epithelium, such as a thick, vascularized follicle wall with dense microvilli and specialized cytoplasmic organelles are common features in the maternal poeciliid placenta, much remains unknown about the ontogeny of the poeciliid follicular placenta (Turner 1940; Grove and Wourms 1991).

To define the genetic components contributing to placental function and examine the selective forces influencing the evolution of this unique poeciliid fish lineage, we constructed a new reference transcriptome for the placental fish *Poeciliopsis prolifica*, the blackstripe livebearer. A placental tissue-specific transcriptome profile was generated by comparison to non-placental tissues from *P. prolifica*, while patterns of protein evolution were compared with other closely and distantly related fish species. *P. prolifica* is a highly matrotrophic poeciliid fish that shares a hypothesized lecithotrophic common ancestor with recently diverged lecithotrophic sister taxa (Reznick *et al*. 2002), thus presenting a model system for examining evolutionary genetic changes proximal to the emergence of the placenta. Notably, we find evidence indicating genetic parallelism, both in function and evolution, of the fish placenta and the mammalian placenta.

## METHODS AND MATERIALS

### Samples

Tissue samples were harvested according to an IACUC approved protocol from captive populations of *Poeciliopsis prolifica* raised at the University of Connecticut. Original stocks were obtained from stock populations at the University of California-Riverside under care of Dr. David Reznick and from Ron Davis, a live-bearer hobbyist in Florida. Both populations originated from the same sample population from the Rio El Padillo in Mexico. Tissues were isolated from fish dissected on ice, immediately snap frozen with liquid nitrogen, and stored at - 80° C. For this study, four sample types were isolated: female brain, liver, whole embryo, and the maternal placental/ovarian tissue complex (MPC). Whole female brain was dissected from the skull and is inclusive of the olfactory bulb, cerebrum, optic lobe, cerebellum and medulla oblongata (to the tip of the spinal cord). Due to its delicate nature, maternal placental tissue was isolated by dissecting whole ovary from pregnant females, excising any fertilized and observable unfertilized eggs, tearing open ovarian follicles, removing developing embryos from those follicles, and reserving the remaining maternal placental/ovarian tissue complex (MPC) that included both ovarian follicles and some remaining ovarian tissue (Figure S1). Late-stage (i.e. nearly full-term) whole embryos, identified by full pigmentation, large size, an ability to persist after being excised from ovarian follicle, and being “late-eyed” (Stage 5 as described by (Reznick 1981)) were sampled and stored with belly sacs intact.

### Sequencing

Two types of sequencing platforms, Roche 454 and ABI SOLiD, were implemented in this study. For 454 sequencing, RNA was isolated from 20 different individuals by homogenizing and disrupting selected tissue samples with syringes in a Trizol solution. Due to individual isolation yields, required template inputs for library construction, and to compensate for among-individual variation, each RNA sample was then pooled by tissue type and mRNA was isolated from 5-10 μg of total RNA using the Poly(A) Purist kit (Ambion). All RNA samples were assessed for quality on a Bio-Rad Experion both pre- and post-Poly(A) extraction. Sequencing libraries were made following standard RNA-Seq library construction protocol for 454 sequencing and sequenced on a Roche 454 Sequencer. To generate SOLiD sequencing data, tissues for three individual MPCs and an embryo from one of these same females were first stored in RNALater and then at -80° C. RNA was isolated by disruption and homogenization of tissues using a Polytron and the RNAeasy mini kit (Qiagen). DNA was removed from each sample by TurboDNAse (Ambion) and validated for sample integrity using an Agilent Bioanalyzer. ERCC spike-in controls (Life Technologies) were then added to each sample and ribosomal RNA (rRNA) was removed using the Ribozero kit (Epicenter). Final RNA-Seq libraries were constructed from the resultant mRNA sample using standard SOLiD transcriptome library construction protocols. Libraries were sequenced on an ABI SOLiD 5500xl.

### Assembly

Post-sequencing, all 454 reads were trimmed using 454 Newbler software to remove bar codes and the program CUTADAPT v1.2.1 (Martin 2011) to remove adapter sequences and trim low quality regions of reads. Seqclean was then used to remove poly-A tails. CUTADAPT was also used for trimming out all barcode and adapter sequences as well as quality trimming for SOLiD libraries. All SOLiD libraries were then screened against an in-house database of rRNA sequences to remove any rRNA sequences that may have not been removed in the rRNA-depletion step. All remaining SOLiD reads were normalized using the Trinity-associated *in silico* k-mer normalization protocols. All trimmed 454 reads and normalized SOLiD reads from all tissues were then input into the Trinity transcriptome assembler (release 7/17/2014) (Grabherr *et al*. 2011). Following the Trinotate pipeline (release 4/30/2015) for annotating predicted transcripts (Haas *et al*. 2013), open-reading frames (ORFs) were predicted using Transdecoder (release 1/27/2015). All transcripts and predicted proteins were then annotated via homology against the SwissProt/Uniprot database and assigned any associated Gene Ontology (GO) terms and eggNOG orthologs group membership. Predicted proteins were also searched for PFAM protein domain and identification as a signaling protein using SignalP (v4.1) (Nielsen 2017), transmembrane protein using TMHMM (v2.0) (Krogh *et al*. 2001), or ribosomal RNA using RNAmmer (v1.2) (Lagesen *et al*. 2007). All transcripts were examined for any additional homologies against the NCBI *nr* database using BLASTX and annotated using BLAST2GO (v2.5.0) (Conesa *et al*. 2005). Any transcript without an *nr* BLASTX-hit was also searched against the NCBI *nt* database with BLASTN. Finally, all transcripts were assessed with BLASTN for homology with known non-coding RNAs (ncRNAs) identified in zebrafish (*Danio rerio*) (Ulitsky *et al*. 2011). Databases versions for all homology searches were all updated on 7/1/15 before this analysis was completed.

Tissue-specific gene expression patterns were surveyed by mapping reads to the Trinity assembled transcriptome sequence, quantifying read coverage among transcripts, and testing for differences among comparison groups. Mapping was performed using BWA (v0.7.7)(SW algorithm) (Li and Durbin 2010) for all 454 data, and Bowtie2 (v4.1.2) (Langmead and Salzberg 2012) for all SOLiD data. Gene expression and read counts were estimated for all transcripts using the program eXpress v1.5.1 (Roberts and Pachter 2013). Count data from 454 mapping was passed through R-based DESeq2 analysis (Love *et al*. 2014) to assess significant differences in pairwise comparisons of gene expression patterns among tissue samples, while correcting p-values for False Discovery Rates (FDR) due to multiple comparison tests. Since sequencing libraries were generated from pooled samples, they were assumed to represent an “average” perspective. Due to the lack of replicates of pooled samples, best practices outlined in the DESeq2 manual were used to generate dispersion estimates by comparing counts among tissue types as opposed to between replicates. This process should be conservative with respect to false positives since it errs on the side of using larger than necessary dispersion values. FPKM (fragments per kilobase per millions reads) values were then used in BioLayout Express3D (v3.2) (Theocharidis *et al*. 2009), along with the MCL (v12-068) clustering algorithm (van Dongen and Abreu-Goodger 2012), to generate a preliminary 3-D gene atlas of co-expressed genes clusters. Due to modest read coverage of 454 sequencing libraries, only “highly” expressed genes (an FPKM value > 50 in at least one tissue) were included in clustering analyses.

### Evolutionary Rates

Evidence of positive selection in the evolutionary rates of poeciliid genes was tested using the branch-sites models implemented in the program PAML v4.7 (Yang 2007). cDNA resources for six other species of fish whose genome and gene models have already been described were downloaded from ENSEMBL and compared to our sequences for *P. prolifica*. These species included the following: *Danio rerio*, *Gadus morhua*, *Takifugu rubripes*, *Oreochromis niloticus*. *Gasterosteus aculeatus*, and *Xiphophorus maculatus* (Figure S2). Of these six species, *X. maculatus* is the most closely-related species to *P. prolifica*; both are in the family Poeciliidae. However, *X. maculatus* differs significantly from *P. prolifica* in reproductive-style since it is a lecithotrophic (yolk-feeding) live-bearer with no evidence of post-fertilization maternal provisioning. *P. prolifica* is highly matrotrophic, with sufficient post-fertilization maternal provisioning to sustain an eight fold increase in dry mass between the fertilization of the egg and birth (Pires *et al*. 2007). Predicted coding sequence regions for *P. prolifica* were compared to cDNA reference sequences for each species using reciprocal best BLAST hit approaches (TBLASTX in this case) to identify orthologous genes between species. Once orthologs were identified, all orthologous gene clusters that lacked a predicted ortholog for any species (i.e. no reciprocal best BLAST hit found) or, when examining high-scoring segment pair (HSP) alignment regions, that yielded a multiple sequence alignments less than <200 bp long were discarded. Using in-house Python scripts, the remaining orthologs were passed through a series of analysis steps. Groups of orthologs were first reconstructed in the same strand and aligned using the codon-guided multiple sequence alignment (MSA) algorithm MACSE v 0.9b1 (Ranwez *et al*. 2011). MSAs were cleaned using trimAl (Capella-Gutiérrez *et al*. 2009) to remove all gaps both from within, and at the ends of, the aligned sequences. MACSE includes the convenient feature of assessing frameshift and stop codon issues associated with multiple sequence alignment. Thus, in order to avoid confounding alignment problems related to poor data quality, low scoring MSAs and true pseudogenized gene sequences, all of which would contribute to false positives in subsequent PAML analyses, this feature was leveraged to identify and remove any MSA with either a frameshift ambiguity or base ambiguity from further analysis.

The remaining MSAs were then analyzed in PAML with three different phylogenetic “foregrounds” to test for positive selection in rapid codon evolutionary rates: *P. prolifica* only, *X. maculatus* only, and all poeciliids. These three levels of examination provided a proxy test of the evolutionary changes possibly associated with three reproductive-styles, respectively: matrotrophic viviparity, lecithotrophic vivparity, and vivparity (generally). Classification of sites having significant evidence for being under positive selection required a significantly better fit of the branch-sites alternative model of positive selection over the null model (implemented as described in the PAML manual – Model 2A vs. Model 1A – with a χ^2^ test using p-value < 0.05 as the threshold for identifying significant improvements in maximum likelihood model fit) and identification using the Bayes empirical Bayes (BEB) method (p-value >0.95). All sites and predicted proteins were compared among different “foreground” analyses to classify protein evolution associated with the aforementioned reproductive-style that these species represent.

A distance-based gene family tree for the *RAB11 family-interacting protein* gene family (*RAB11FIP*) was constructed using neighbor-joining tree methods to describe the general patterns of gene duplication and evolution in fishes. Jukes-Cantor distances among protein sequences were used to generate tree topology. All sequences included in this gene family tree where gathered by identifying any *P. prolifica* predicted protein sequence with homology to *RAB11FIP*s in *Danio rerio* using BLASTP (e-value < 1e-5) and using those predicted proteins to identify any other existing protein sequences for *RAB11FIP* genes in fishes using BLASTP (e-value <1e-5; taxonomically restricted search to “bony fishes” – taxid: 7898). MUSCLE v3.8.31 (Edgar 2004) was used to generate a multiple sequence alignment for all sequences and CLC Genomics Workbench v7.5 was used to generate a tree with 100 bootstraps. To focus analysis on *RAB11FIP* genes only, all clusters of genes identified as the protein *UNC-13* (a homologous gene to *RAB11FIP*s) were trimmed from final tree.

### Data Availability

All read data was deposited in the NCBI SRA database under the following accession numbers: SRR1639275, SRR1640127, SRR1640137, SRR1640160, SRR1640171, SRR1640200, SRR1640209, SRR1640216, and SRR1640219 under the BioProject PRJNA266248. All custom scripts are available here: https://github.com/juefish/Jue_et_al_G3_P_prolifica_transcriptome.git.

## RESULTS

### Assembly Statistics

*De novo* assembly of 3,696,154 Roche 454 and 159,802,508 SOLiD reads (post-trimming, see Table S1 for library details) yielded a transcriptome of 331,767,677 Mb (43.74% GC) with 478,065 predicted transcripts (TSA Reference ID: GBYX00000000.1). Average contig length was 639 bp and N50 was 885 bp. These contigs were grouped into 319,532 components, which are analogous to estimated “genes” or groups of isoforms (Table 1). While some of these predicted transcripts could be spurious or fragmented results from the assembler, 236,360 (49.4%) of these predicted transcripts were well-supported with read depth of coverage >10x, representing a very diverse transcriptome (Table 1).

Within this assembled transcriptome, 113,240 transcripts (23.6% of total) were predicted to have a protein open-reading frame (ORF) (Figure 1), with over 80% of these predicted proteins (both total transcripts and genes) carrying homology with a protein in the UniProtKB/Swiss-Prot database, and >75% of those showing associations with known Pfam domains (Figure 1). Functional Gene Ontology (GO) annotations were identified for the majority of these sequences with homology to *nr* database reference sequences, representing a multitude of functional elements, spanning a range of categories in the Gene Ontology (Figure 2). Another 41,851 transcripts with no BLAST result at all (8.7%) showed similarity to REPBASE repetitive element sequences, including 1,747 transcripts from 1,043 predicted genes that incorporated repetitive element genes (Table S3). These transcripts span a wide-range of repetitive element origins, including elements known to have specific placental function in mammals such as *retrotransposon-derived protein PEG10-like*. Another 286 transcripts carry regions identified by homology with non-coding RNAs from *D. rerio*. These transcripts represent a variety of non-coding RNAs that may be involved in gene regulation (Table S4). For instance, one identified transcript shows homology with *cyrano*, a lncRNA demonstrated to be necessary for proper embryonic development and interacting with a known miRNA miR-7 (Ulitsky *et al*. 2011). A small number (24) of these transcripts showed evidence for bidirectional transcription and, thus, candidates for active functioning in gene regulation through complementary base-pairing with coding transcripts.

**Figure 1.**
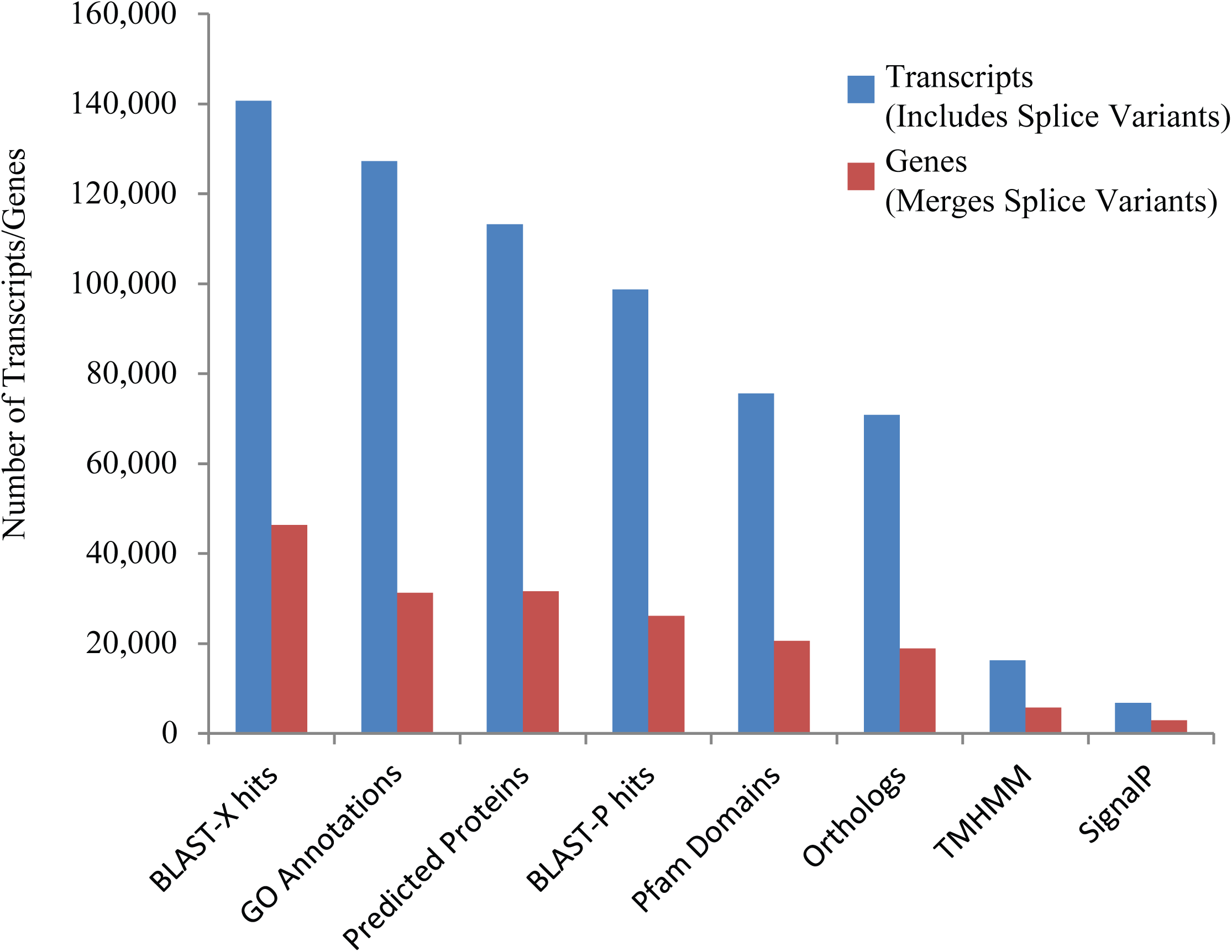
Distributions of various transcriptome annotations for *Poeciliopsis prolifica* reference transcriptome predicted transcripts (blue) and alternatively-spliced variant groups, representing “genes” (red). 140,709 transcripts (29.4% of total) exhibited identifiable homology (e-value < 1 x 10^-5^) with protein reference sequences in the NCBI *nr* database and another 29,199 (6.1%) transcripts showed similarity (e-value < 1 x 10^-5^) with nucleotide reference sequence in the NCBI *nt* database. 16,277 (11.6%) and 6,772 (4.8%) transcripts are associated with transmembrane (TMHMM) and signaling (SignalP) proteins. 8,181 showed greater than 70% coverage of known UniProtKB/Swiss-Prot orthologs; 3,785 transcripts were identified as containing the complete ORFs of conserved orthologs in UniProtKB/Swiss-Prot database (Table S2).

**Figure 2.**
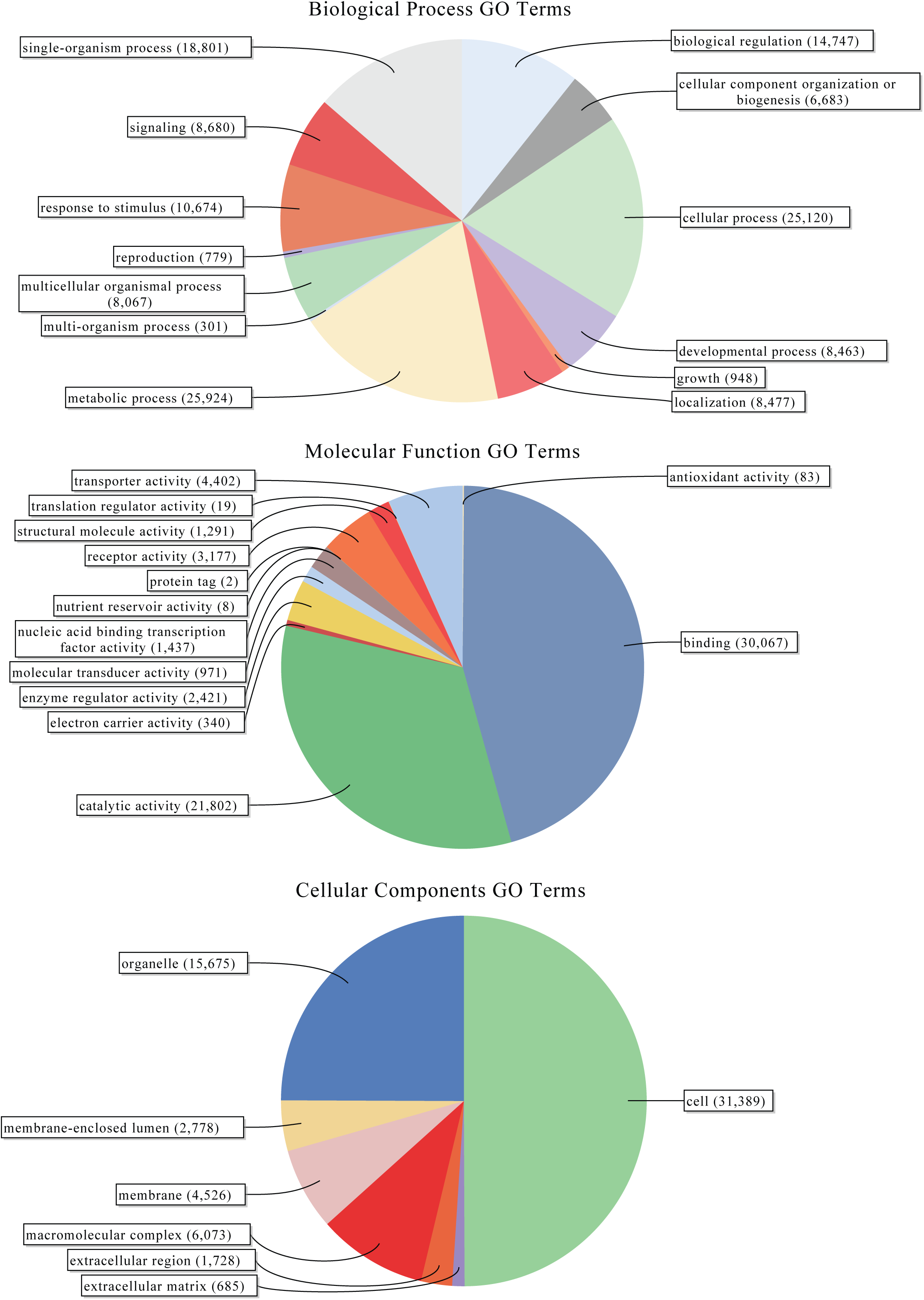
Level 2 gene ontology term distributions for reference transcriptome of *Poeciliopsis prolifica*.

### Tissue Specific Gene Expression

Using MCL clustering of gene expression estimates, we generated a preliminary gene atlas for *P. prolifica* to identify clusters of co-expressed transcripts among four different sample types: MPC, female brain, liver, and late-stage developing embryo, hereafter referred to as “tissues”. Before clustering, pairwise tests for significant differences (p-value <0.05 after correction for FDR) in gene expression using DESeq2 were conducted across all transcripts in all tissues and revealed 45, 108, 18, and 24 transcripts were specifically expressed in MPC, whole embryo, brain and liver, respectively. For MCL clustering analysis and gene atlas construction, a subsample of the 6,839 most highly expressed transcripts (FPKM values >50 in at least one of the four tissues) were included in the analysis. This subset further reduced the number of identifiable (via pairwise comparisons) tissue-specific transcripts included in the atlas that were significant for tissue-specific expression to 24, 36, 4, and 5 for MPC, embryo, brain and liver, respectively. Using the tissue-specific gene expression patterns of these transcripts (Figure S3) and the MCL clustering algorithm, nine co-expressed gene clusters were identified (Figure 3).

**Figure 3.**
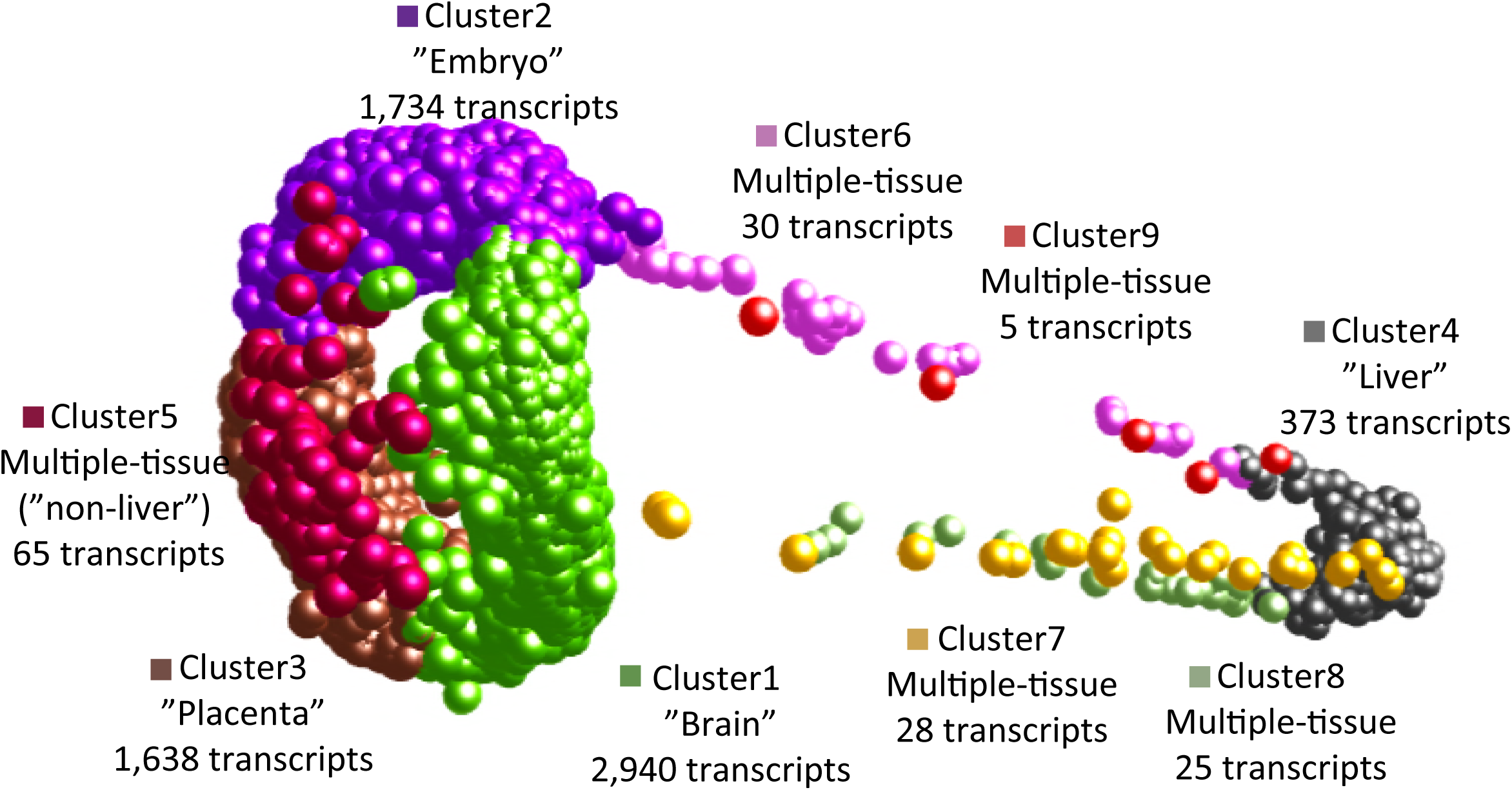
Three-dimensional gene atlas derived from gene expression data for maternal placental/ovarian complex (MPC), late-stage embryonic, brain, and liver tissue. Proximity in space indicates similarity in gene expression profile across tissues. Clusters were defined using MCL clustering algorithm on highly expressed genes (>50 FPKM in at least on tissue type) from Roche 454 RNA-seq. Clusters 1-4 are mostly, though not exclusively, made up of transcripts that are tissue-specifically expressed, while clusters 5-9 consist of transcripts that are highly expressed across multiple tissues. Each of these clusters (1-9), had 2,940, 1,734, 1,638, 373, 65, 30, 28, 25, and 5 members, respectively.

Cluster 1 was the largest cluster and generally associated with transcripts that have high expression in the brain, but showing some co-expression with other tissues, particularly MPC and embryo. Cluster 2 was generally associated with transcripts highly expressed in embryo, cluster 3 was associated with transcripts highly expressed in MPC, and cluster 4 was associated with transcripts highly expressed in liver. Clusters 5 to 9 (which represented only 2.2% of the transcripts in the atlas) were defined by expression across multiple tissue types, displaying gene expression profiles indicative of “house-keeping”-like genes (Figure S3). Transcripts with significant evidence for tissue-specific expression largely supported these cluster classifications with 32 of the 36 aforementioned “embryo”-specific genes in cluster 2 and all 24 of the MPC genes in cluster 3. Brain and liver clusters were less clearly supported with none of the four “brain” genes in cluster 1 and only one of the five “liver” genes in cluster 4; however, the number of transcripts in these clusters was so low that detectability may have been limited.

Transcripts involved in progesterone signaling pathways were observed as highly expressed in placental tissues. Overall, 242 transcripts with ORFs were identified as having GO-associations with progesterone regulatory pathways, including *Protein DEPP* (*decidual protein induced by progesterone*), suggesting that similar developmental patterns in cell differentiation and specialization maybe be occurring in fish as it does in mammals during pregnancy (Watanabe, et al. 2005).

### Repetitive Element Transcripts

Repetitive element gene expression was observed across various tissue samples and a subset of the gene atlas clusters. Of the clustered 454 expression data, the MPC cluster (#3) had the highest number of repetitive element transcripts, with a total of 9 transcripts; the “brain” cluster (#1) had the second highest repetitive element transcript count at five transcripts. Cluster 2 (embryo), cluster 4 (liver), and cluster 5 (multiple tissues) had 2, 1, and 1 transcript(s), respectively. Only one transcript of these 18 transcripts found in the gene atlas clusters (identified as a *transposable element tc1 transposase*) showed no expression in placenta; all 17 other transcripts were expressed (>50 FPKM) in MPC (eight of these transcripts were also identified as homologs to *transposable element tc1 transposases*). One transcript (a *reverse transcriptase*) was also identified using the aforementioned pairwise significance testing (DEseq2, p-value < 0.05) as more expressed in MPC as opposed to other tissues (FPKM _MPC_ = 155.7 vs. FPKM _average_other_tissues_ = 5.07). The three MPC SOLiD libraries also indicated high levels of MPC gene expression of repetitive element-derived transcripts. From the SOLiD RNA-Seq data, 98% of the 1,747 transcripts from the broader transcriptome reference sequence and originating from repetitive elements were expressed in either MPC or embryonic tissues, with 227 predicted transcripts from 199 predicted genes expressed either only in the MPC or >5 fold greater expression in MPC over embryonic tissues (Table S3). Approximately an equal number, 213 predicted transcripts and 199 predicted genes were found associated with embryonic tissues using the same criteria (Table S3). Eight transcripts had an FPKM value of >50 across and were identified as four gene families that included an envelope protein, a partial pol protein, a tc1 transposase and a tc3 element. In addition to the gene classes mentioned above, other repetitive-element transcripts were identified as *retrotransposon-derived protein PEG10-like*, *120.7 kDa protein in NOF-FB transposable element*, *retroviral polyprotein*, and *transposable element tcb1 transposases*. These transcripts appeared unique to the poeciliid lineage, showing between 50% and 70% similarity to other repetitive element reference sequences from other species, with only *retrotransposon-derived protein PEG10-like* showing high similarity (88%) with reference sequences from the NCBI *nr* database.

### Transcripts with Unknown Function

The majority of these clusters of highly expressed genes consisted of transcripts with no known annotation. Of the highly expressed transcripts described in these clusters, 79.4% (n=6260) were not identifiable via BLAST searches of SwissProt/UniProt, *nr* and *nt* databases (e-value < 1 x 10^-5^). A large number (786, or 12.6%, of the total unknowns) of these predicted transcripts had evidence for some type of repeat in their sequence, with 761 of the repeats identified as either a simple repeat or low complexity sequence, indicating that the sequence may be part of a non-coding region (Wren *et al*. 2000; Morgante *et al*. 2002; Liu *et al*. 2012). Many of these sequences are likely either species-specific 5’ or 3’ UTRs or previously undescribed non-coding RNAs. For example, another four of these transcripts in this cluster were associated with known non-coding RNA sequence from *D. rerio* (3 with miRNAs and 1 with a lncRNA); however, given that all of these sequences were much longer than miRNA size (312-982 bp) and not readily identifiable as miRNA precursors (Liu *et al*. 2015), they are more likely to be binding sites for such targets than host transcripts. Another 49 transcripts had predicted ORFs associated with them, but no BLAST annotation and thus appear to be novel protein sequences. Of these 49 predicted proteins, two were identified as prospective signaling peptides, one of which was a member of the MPC gene cluster. The other “signaling” peptide and two other predicted proteins were identified as transmembrane proteins. The signaling/transmembrane protein was a member of the “house-keeping gene” cluster (but most highly expressed in liver), while the other two transmembrane proteins were associated with either the “brain” cluster or the “embryo” cluster. Notably, the “embryo” cluster member was also highly expressed in MPC (FPKM_embryo_=53.5; FPKM_placenta_=39.5). Given exhaustive attempts to annotate these sequences and the fact that they are highly expressed transcripts, these sequences appear to be novel to this species.

### Protein Evolutionary Rates

Reciprocal best BLAST hits of the cDNA coding sequence against the predicted and known cDNAs for six fish species with sequenced genomes revealed predicted *P. prolifica* transcripts to have 12,631 orthologs with *Danio rerio*, 14,761 orthologs with *Xiphophorus maculatus*, 12,899 orthologs with *Takifugu rubripes*, 12,316 orthologs with *Gadus morhua*, 13,388 orthologs with *Gasterosteus aculeatus*, and 13,282 orthologs with *Oreochromis niloticus*. Out of all of these orthologs, only 5,398 were shared orthologs for all seven species (including *P. prolifica*). Within this shared ortholog set, 963 ortholog alignments showed evidence of open-reading frame indels in at least one species’ orthologous sequence, resulting in a frame-shift in predicted codon sequences (Table S5). These frame-shifts could be the result of errors in a given fish reference sequence or bona fide mutations in a specific species. While all species showed evidence for frame-shifts, transcript sequences from *D. rerio*, *P. prolifica*, and *X. maculatus* had a higher proportion of orthologs with an identified frame-shift than the remaining species (Table S5). Additionally, 978 ortholog groups were discarded from the PAML analysis due to ambiguous bases (“N”) in the reference sequences; this was a disproportionately acute issue with *G. morhua* sequences (912 orthologs).

Within the final set of 3,457 orthologs employed in our PAML analyses, 2,298 sites across 404 predicted proteins were identified as undergoing positive selection. Of these sites, 917, 1104, and 247 were associated with *P. prolifica*, *X. maculatus*, and both poeciliids, respectively (Figure 4, Table S6-S13). The predicted proteins carrying these sites covered a wide-range of biological functions (Figure S4) with no overall significant enrichment for any specific functional GO terms relative to the overall transcriptome annotation. Comparisons between the matrotrophic *P. prolifica* and lecithotrophic *X. maculatus* orthologs with sites under positive selection showed genes under positive selection in *P. prolifica* to be significantly enriched for a variety of GO terms over those found in *X. maculatus* (Figure 5, FDR p-value < 0.05). The terms were generally associated with Biological Processes related to biosynthesis and regulatory processes, Molecular Functions terms related to nucleic acid binding, and Cellular Components terms related to the nucleus. Of these sites, 1,376 occurred in regions of these open-reading frames that carried no discernable, previously known protein domain defined by Pfam database searches. Thus, these sites indicate possible novel functional domains for these proteins in *P. prolifica*.

**Figure 4.**
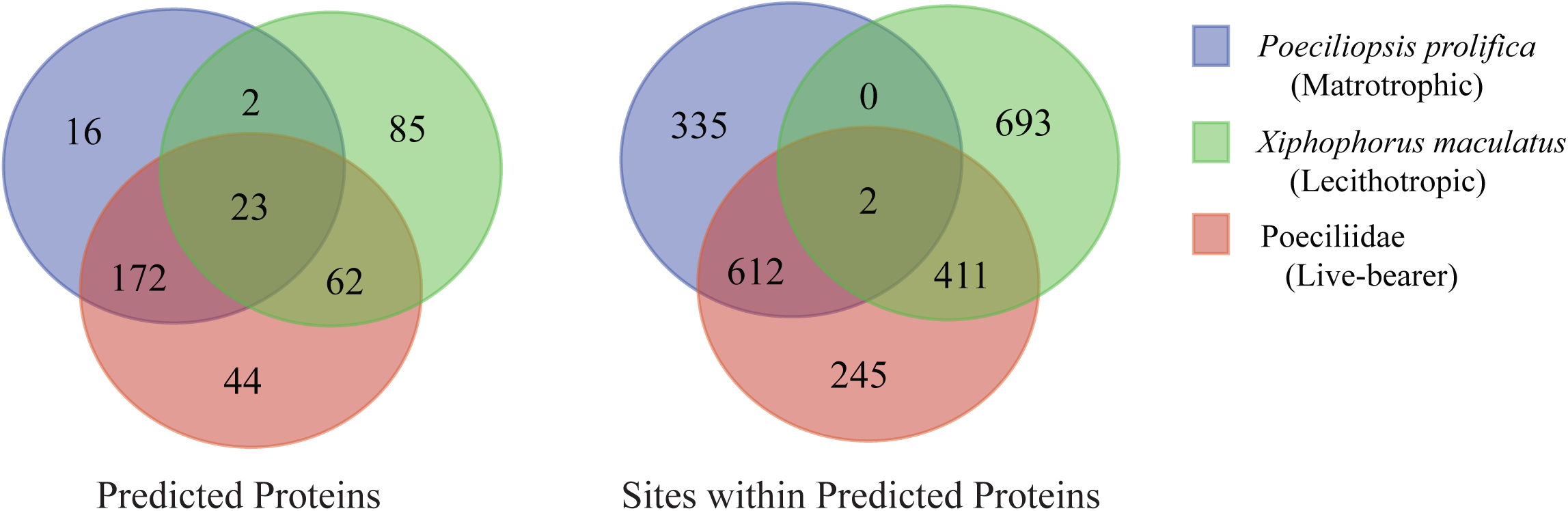
Venn diagrams showing patterns of shared and unshared proteins and sites within protein under positive selection among the three foreground taxon groupings tested with PAML.

**Figure 5.**
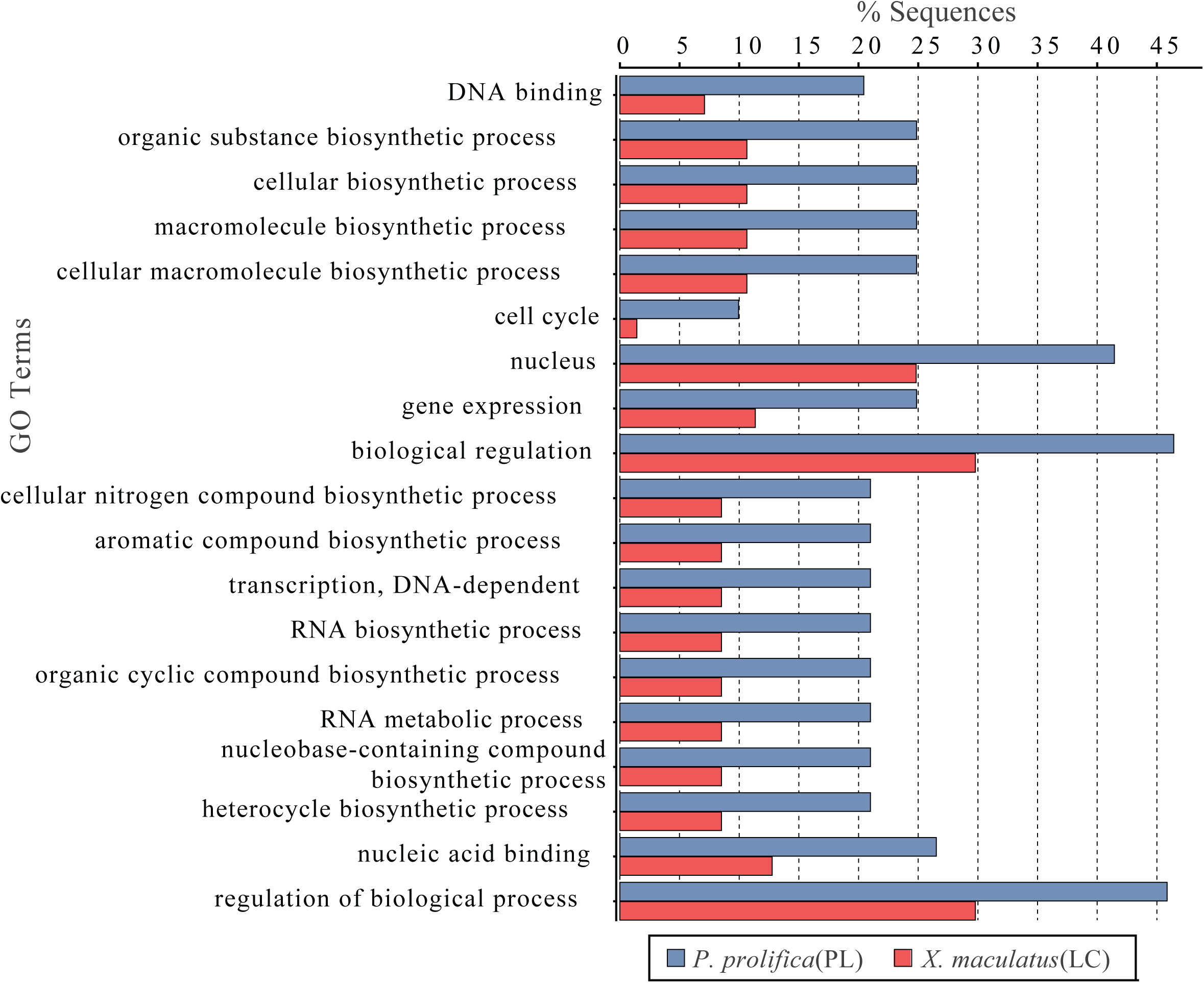
Distribution of GO Terms that were differentially represented in genes identified to be under positive selection in the matrotrophic/placental(PL) *Poeciliopsis prolifica* and lecithotrophic(LC) *Xiphophorus maculatus*. GO terms include categories from all three main ontologies (Biological Processes; Molecular Functions; Cellular Components).

While the majority of proteins undergoing positive selection (67%) had less than five sites identified as under positive selection, many of the genes under positive selection exhibited evidence for extensive rapid evolution (Table S7). For instance, the *GRAM domain-containing protein 4*, *GRAMD4*, carries 94 sites identified as evolving rapidly in *X. maculatus*. These sites account for 16% of the entire protein sequence for this gene. None of these sites overlap with the known GRAM protein domain, indicating that this region may be an important novel functional domain. *GRAMD4* is a membrane protein known to be a tumor suppressor in apoptotic pathways associated with mitochondria (John *et al*. 2011). *Insulin-like growth factor 1a receptor* (*IGF1RA*) is another gene that has a large number of sites under positive selection in *X. maculatus*. Overall, 96 sites within *IGF1RA* were shown to be under positive selection, with eight sites showing changes in both poeciliids, while the remaining 88 were restricted to *X. maculatus* (Figure 6).

**Figure 6.**
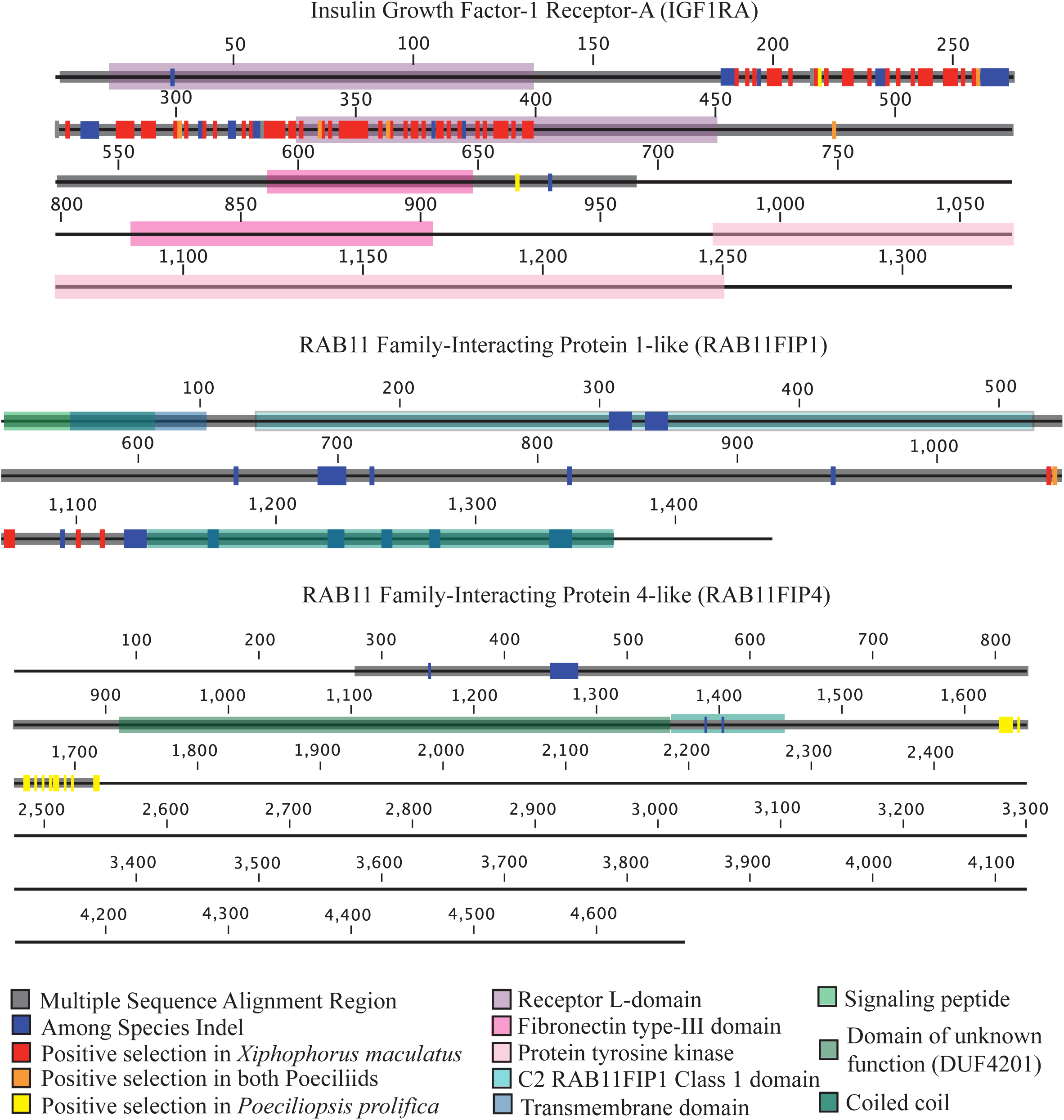
Diagrams of *insulin growth factor-1 receptor-A* (*IGF1RA*) from *Xiphophorus maculatus*, and *RAB11 family-interacting protein 1-lik*e and *4-like* (*RAB11FIP1* and *RAB11FIP4*, respectively) from *Poeciliopsis prolifica* showing known protein domains, indel regions among species (identified using regions of multiple sequence alignment), and sites identified as being under positive selection from PAML analysis in live-bearing poeciliids, the lecithotrophic *Xiphophorus maculatus*, or the matrotrophic *Poeciliopsis prolifica*.

Protein lengths for *IGF1RA* vary among species. In our *P. prolifica* assembly, we have predicted only 711 residues for this protein, but our sequence may be incomplete as it lacks a 3’ UTR region. Within *X. maculatus* where there is a complete predicted gene sequence (1,332 aa), these 96 sites account for ∼7% of the gene sequence. Of the 96 sites, 48 are located within the Furin-like domain of the protein, 36 are in one of the Receptor L-domains, one is in the Fibronectin type III domain, and 11 are found outside of any known protein domain.

*P. prolifica* generally showed different genes under positive selection than *X. maculatus* (Figure 4; only 17.1% of the 404 orthologs under positive selection showed positive selection in both species). For example, *RAB11 family-interacting protein 4-like* (*RAB11FIP4*), one of the six types of *RAB11 family-interacting proteins* found in fishes (Figure S5), has 16 sites under positive selection in *P. prolifica*, but none in *X. maculatus*, while another member of that same gene family, *RAB11 family-interacting protein 1-like* (*RAB11FIP1*), has 5 sites under positive selection in *X. maculatus* and 1 in both *X. maculatus* and *P. prolifica* (the 2 species have different residues at that site). These sites may be associated with novel functional domains because each of these sites were identified as being extracellular for both *RAB11FIP1* and *RAB11FIP4* using the transmembrane identification algorithm TMHMM; however, none of these sites are located within any “known” functional domain (Figure 6). Patterns of gene evolution across fish species show that the rapid gene evolution may be likely facilitated by multiple incidences of gene duplication. Along with *IGF1RA*, *RAB11FIP* gene family members showed family-wide evidence for gene duplication events and both *RAB11FIP* genes that were shown to be under positive selection had expressed paralogs in the reference transcriptome sequence (Figure S5). These duplications likely occurred after the whole genome duplication event experienced by all fishes (Jaillon *et al*. 2004) since there is only one copy of each family member found in the gar, *Lepisosteus oculatus*, (Figure S5) which has not undergone the teleost fish whole genome duplication event.

## DISCUSSION

We have developed the most thorough transcriptome reference for a placental fish to date, providing a significant extension to earlier work in a sister taxa (Panhuis *et al*. 2011), in order to better understand the genetics and evolution of placentation in fish. Our sequence assembly has been extensively annotated for functional content and provides a solid foundation for establishing genomic resources for this genus. Identified transcripts cover diverse functions and, given the sampling of both poly-A selected and ribo-depeleted RNAs across multiple tissues, provide a comprehensive assessment of both protein-coding and non-coding RNA genes organism-wide. In addition to its general descriptive characteristics, this transcriptome reference has also provided us with important insights into the genetics of this placental species.

There appears to be parallels in placental evolution in eutherian mammals and *P. prolifica*, highlighted by the extensive presence of expressed repetitive elements in fish MPC tissues. Eutherian mammals often utilize repetitive element components as functional contributions to placental and embryonic development, including endogenous retroviral envelope proteins (Mi *et al*. 2000), DNA transposon regulatory machinery (Lynch *et al*. 2011), and/or *gag* and *pol* domains of LTRs (Ono *et al*. 2001). A total of 98% of the transcripts associated with retroelements exhibited high expression to either placental or embryonic tissues. These transcripts included a variety of orthologous genes associated with placental function in mammals, such as *PEG10*, an imprinted gene expressed in the placenta of mammals. The extensive presence of progesterone signaling-related genes also parallels mammalian placental function, particularly functions associated with the corpus luteum (Gemmell 1995) and decidual cells (observed expression of *Protein DEPP* in fish MPC parallels that also described in mammalian placental and embryonic tissues (Watanabe *et al*. 2005)). Alternatively, expressed repetitive element transposases may be co-opted genes involved in more general gene regulation as transcription factors or DNA-binding proteins with centromeric functional roles (Feschotte 2008). The identification of seemingly convergent gene expression of genetic elements of similar type, but different lineage and an apparent implication in the function of the independently derived placental tissues of fish and mammals leads to a hypothesis that similar molecular and cellular adaptations are functioning in both systems.

There was also extensive evidence for placental tissue usage of novel genes and transcripts as functional components specific to this family of fishes and, possibly, restricted to this species. Tissue-specific patterns of high gene expression implicate many novel components to be active in the MPC. Most of these novel, predicted transcripts lacked homology to genes in any existing genetic resource, strengthening support for their designation as “novel”. In total, 17.6% of the predicted protein sequences in the reference transcriptome could not be associated with any existing reference sequence via exhaustive comparison to known protein and coding sequence databases. Mis-assembly and/or chimeric reads could only explain a minority of these “unknowns” as the depth of coverage was generally high for these genes and, as evidenced by the clustering analysis, many of these transcripts are highly expressed. Many “unknowns” (∼9%) were found to contain repetitive elements or have sequence homology with non-coding RNAs, implicating the co-option of rapidly evolving elements in the origins of this novel transcriptional diversity. As our pairwise ortholog identification shows, the closer the phylogenetic species comparison is, the greater the proportion of the transcriptome we could identify and annotate (e.g. 14,761 orthologs were found in *X. maculatus* vs. 12,631 orthologs in *D. rerio* for a 16.9% increase in the number of identified orthologs). Overall, novel transcripts would appear to be significant contributors to placental function in *Poeciliopsis*. This prediction is also congruent with mammalian placental systems, wherein many of the transcripts observed associated with placental development and function are derived from lineage-specific co-option and domestication of typically inactive retroelements (Emera and Wagner 2012).

### Role of Protein Evolution/Positive Selection

Using our reference sequence, we identified genes under positive selection in both matrotrophic (P. prolifica) and lecithotrophic (X. maculatus) species of livebearing poeciliid fishes. Overall, genes identified as under positive selection did not disproportionally represent any specific functional group, indicating that any genetic signal of adaptation identified in this analysis covered a wide-array of functional components in the Poeciliidae. However, the statistically significant differences in functional groups among the lecithotrophic *X. maculatus*, and the matrotrophic *P. prolifica* undergoing positive selection indicate that there may be selective bias in the types of genes contributing to the rapid evolution of placentation in this group. The identification of genes related to biosynthesis and gene regulation, especially those associated with DNA-binding in nuclear regions, are significantly over-represented in genes under positive selection in our placental species. That these functional categories would be under strong selective pressure is consistent with the inherent requirement for placental tissues to develop quickly to support embryonic growth as well as the potential for parent-offspring intragenomic conflict. The unexpectedly extensive protein-coding sequence evolution is highly relevant to continued interest in the relative contribution of either changes at the protein-coding level or those in gene regulation contributing to evolutionary patterns (Hoekstra and Coyne 2007; Lynch and Wagner 2008; Stern and Orgogozo 2008).

While the majority of genes under positive selection contain only a few sites that are rapidly evolving, some genes exhibit evidence of surprisingly large regions of their coding sequence under Darwinian selection. Evidence of gene duplication would appear to facilitate the potential for positive selection. For example, while the insulin-like growth factor signaling axis is a key regulator of embryogenesis and fetal growth in all vertebrates (Schlueter *et al*. 2007), there is considerable redundancy in many of its components in fishes due to the ancient WGD (Jaillon *et al*. 2004). Specifically, there are multiple copies of *insulin growth factor receptor 1* (paralogs A and B). It has been established for the genus *Poeciliopsis* that *IGF2* has evolved under positive selection that is hypothesized to be driven by parent-offspring conflict (O’Neill *et al*. 2007). While *IGF2* is excluded in our analysis due to stringency filters (it has a large indel region in *P. prolifica* and the HSP alignment region was too short for inclusion), *IGF1RA* (also known to be expressed in fish gonadal tissues (Mei *et al*. 2014)) was shown to have extensive evidence for rapid evolution in this group with eight sites evolving rapidly in all Poeciliidae and 84 sites under positive selection in just *X. maculatus*. These 92 sites cover both conserved protein domains and unannotated regions of the protein. The signal for positive selection on both *IGF2* and *IGF1RA* in poeciliids may reflect the opposing parent-specific expression (imprinting) of *IGF2* and its antagonistic receptor *IGF2R* in mammals. However, if conflict is driving this pattern, then it seems to be pushed to extremes in the non-placental *X. maculatus*, where *IGF2* has also been shown to be under positive selection (Schartl *et al*. 2013). This would appear to contradict assumptions of the hypothesis that parent-offspring conflict would be more extensive in placental species (Zeh and Zeh 2008); alternatively, this observation may indicate that conflict manifests itself differently in the absence of material exchange between mother and fetus. For instance, selection may be acting on the duration of ovoviviparous development in *X. maculatus*, where different paternal genomes compete for gestational space within the mother, while the maternal genome dictates the length of her pregnancy and maximum occupancy of gestational spaces.

While, speculatively, *IGF1RA* may exhibit evidence for selective pressure due to genetic conflict in the lecithotrophic *X. maculatus*, it is possible that other biological processes specific to viviparous reproduction are also under selection in *P. prolifica*. For example, the *RAB11FIP* genes show lineage-specific patterns of protein evolution, indicating different selection pressures in *P. prolifica* and *X. maculatus*. *RAB11FIP*-associated proteins are typically identified by the presence of a C-terminal Rab-binding domain and are involved in vesicle transport and recycling (Lindsay 2004), protein trafficking and sorting (Peden *et al*. 2004) and recycling of membranes in cytokinesis (Wallace *et al*. 2002). It is unclear precisely why these genes are under positive selection in these fish, but given their defined functions they may be responding to lineage-specific selection pressure involving cellular transport related to the evolution of live-bearing.

Gene duplication is likely providing a considerable contribution to the potential for these genes to undergo changes due to positive selection (e.g. Steinke *et al*. 2006). Just as in the *IGF1R* genes, each of these two *RAB11FIP* genes has a closely related paralogous copy that showed no evidence for positive selection (Figure S5).

Overall, our study demonstrates patterns of both sequence and functional convergence of the poeciliid placenta with the therian mammalian placenta. In contrast to predictions that genetic components would be distinct to the poeciliid lineage given the relatively recent convergent derivation of the fish placenta from the pericardial sac and it highly dissimilar structural form, many of the genetic components that contribute to mammalian placental development and function are also involved in the fish placenta. While it could be predicted that at least some of the types of genes involved in placentation in both lineages would be similar with respect to cellular function and functional requirements of any placenta in maternal-fetal exchange, it is notable that we find parallel evolutionary mechanisms, beyond such cohorts of genes, evident in the co-option of retroelements and gene duplication as key contributors to the evolution of this complex organ.

## Acknowledgements

Thanks to Ron Davis for volunteering fish from his personal stocks to support this work and Valeria Pellicer for her illustration. This work was supported by NSF Project Grant IOS-0920088 to MJO and RJO and NSF-MRI 0821466 to Linda Strausbaugh and RJO, and an MRI-R2 to MJO and RJO. This work was supported by the facilities of the Center for Genome Innovation (formerly the Center for Applied Genetics) within the Institute for Systems Genomics.

---

**Figure S1.**
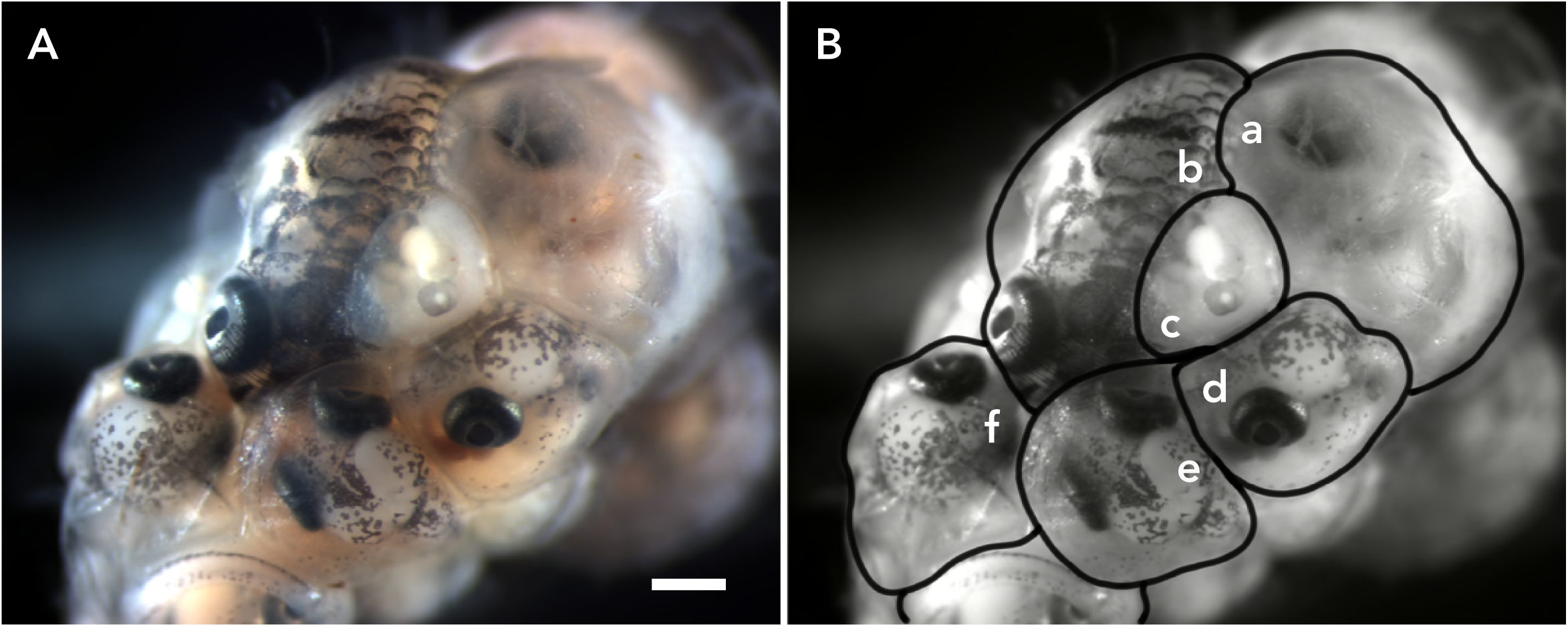
**A and B.** Intact ovary removed from gravid female. Scale bar = 0.5mm. **B.** Outlines of the different embryos within the ovary shown in **A. a.** maternal/placental ovarian tissue complex (MPC) with late stage embryo removed, **b.** Very Late-eyed stage embryo (i.e. nearly full term, Stage 6), **c.** Early-eyed stage embryo ge 3), **d.-f.** Late-eyed stage embryo (Stage 5).

**Figure S2.**
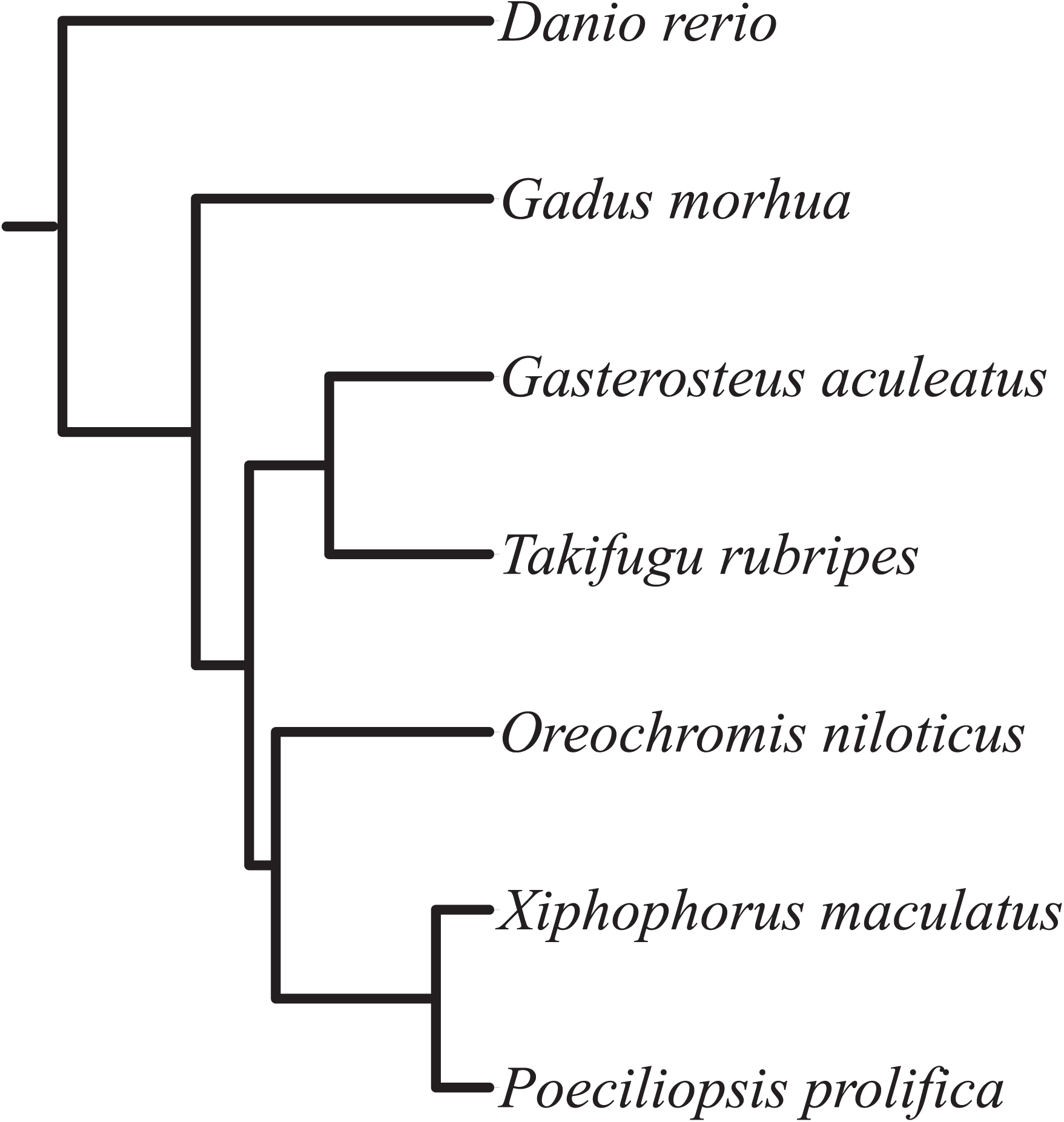
Species tree used in PAML. Tested foregrounds of live-bearing Poeciliids generally, lecithotrophic live-bearers, and matrotrophic live-bearers using the clade of Poeciliids, the *Xiphophorus maculatus* lineage, and the *Poeciliopsis prolifica* lineage, respectively for each case.

**Figure S3.**
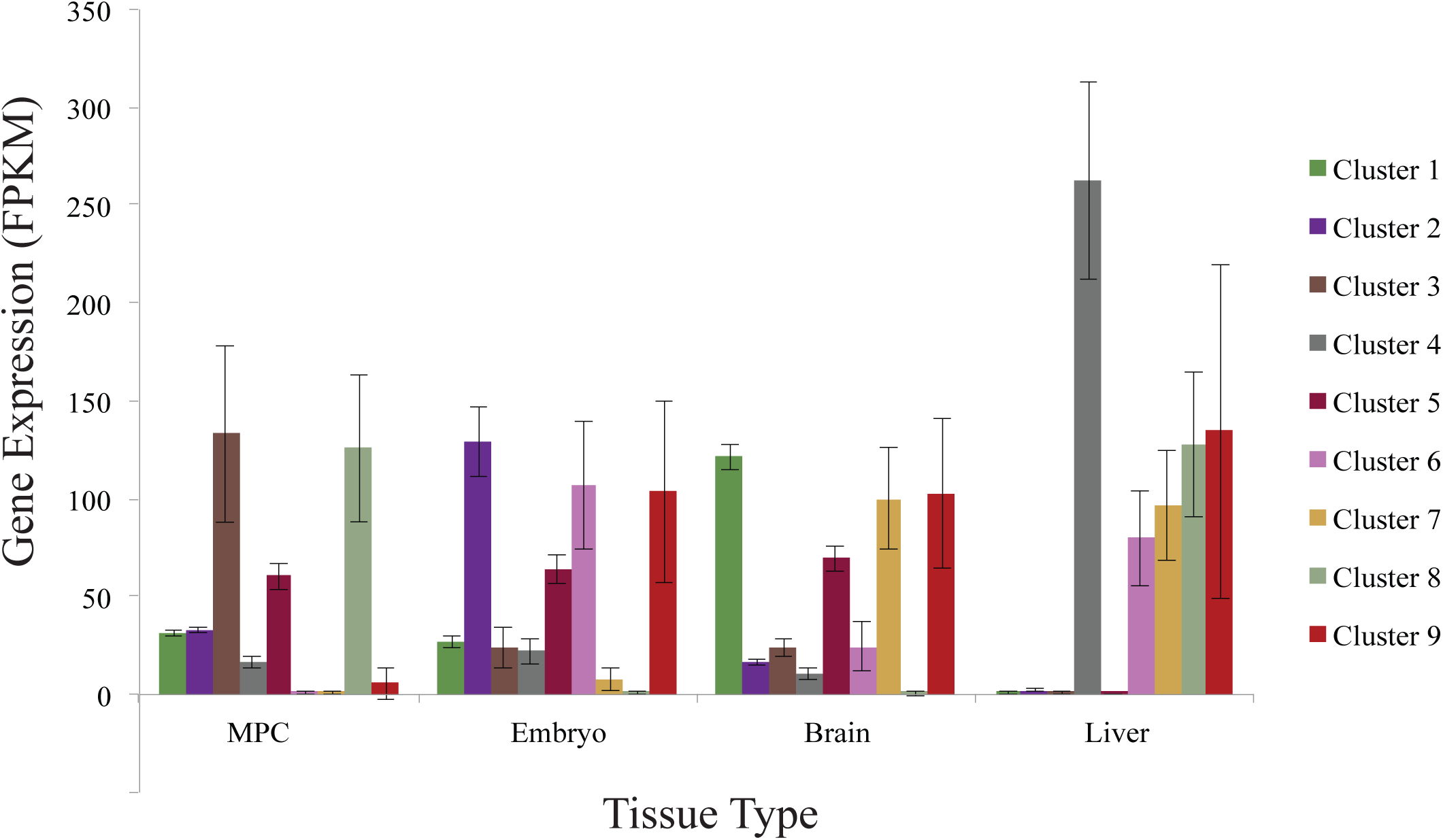
Average expression in fragments per kilobase per million base pairs (FPKM) in each tissue grouped by cluster genes indicating which clusters are associated with which tissues. Standard error bars shown. MPC indicates maternal placental/ovarian complex tissue sample.

**Figure S4.**
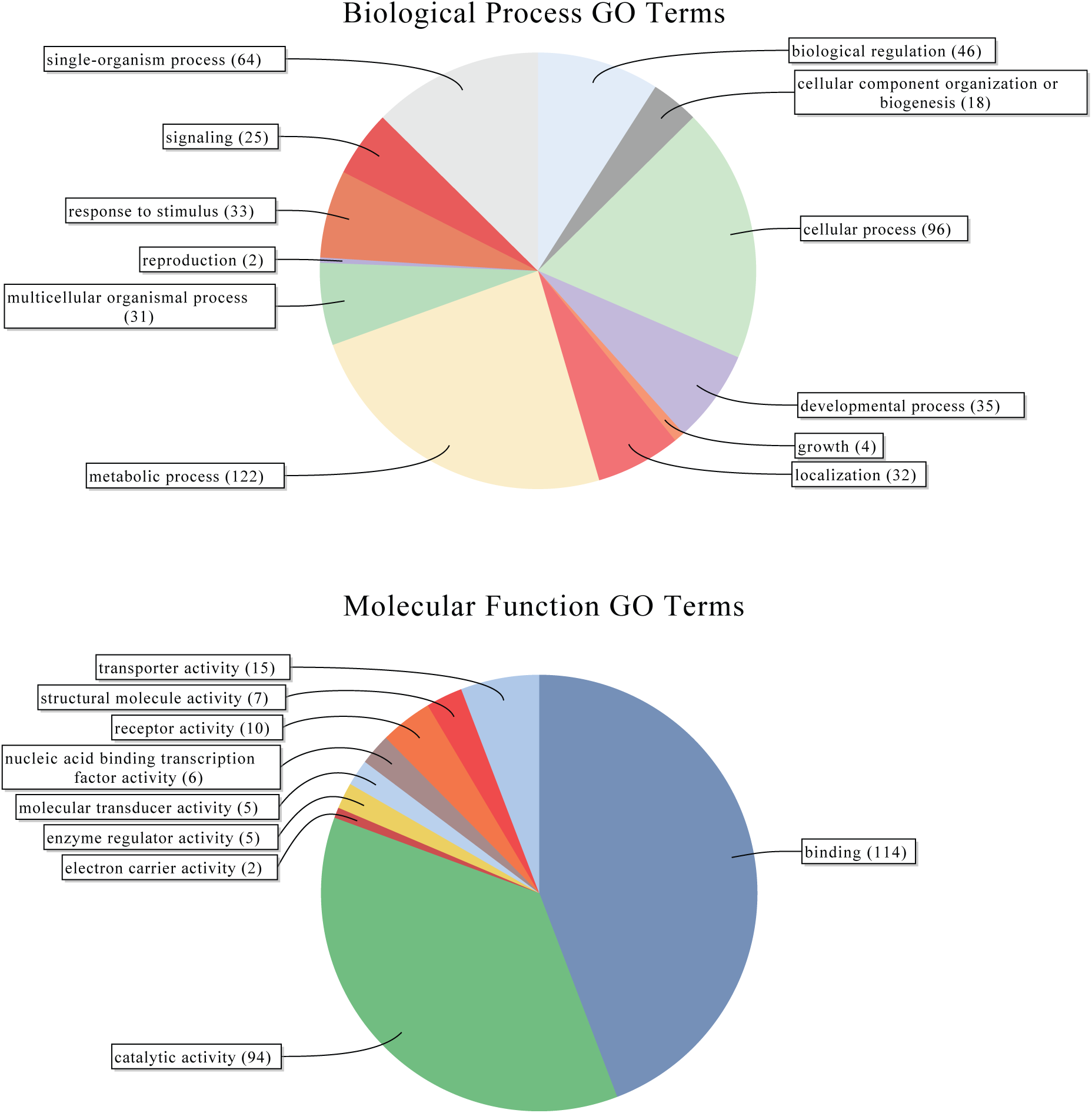
Level 2 Biological Process and Molecular Function Gene Ontology terms associated with gene identias having sites under positive selection in poeciliids.

**Figure S5.**
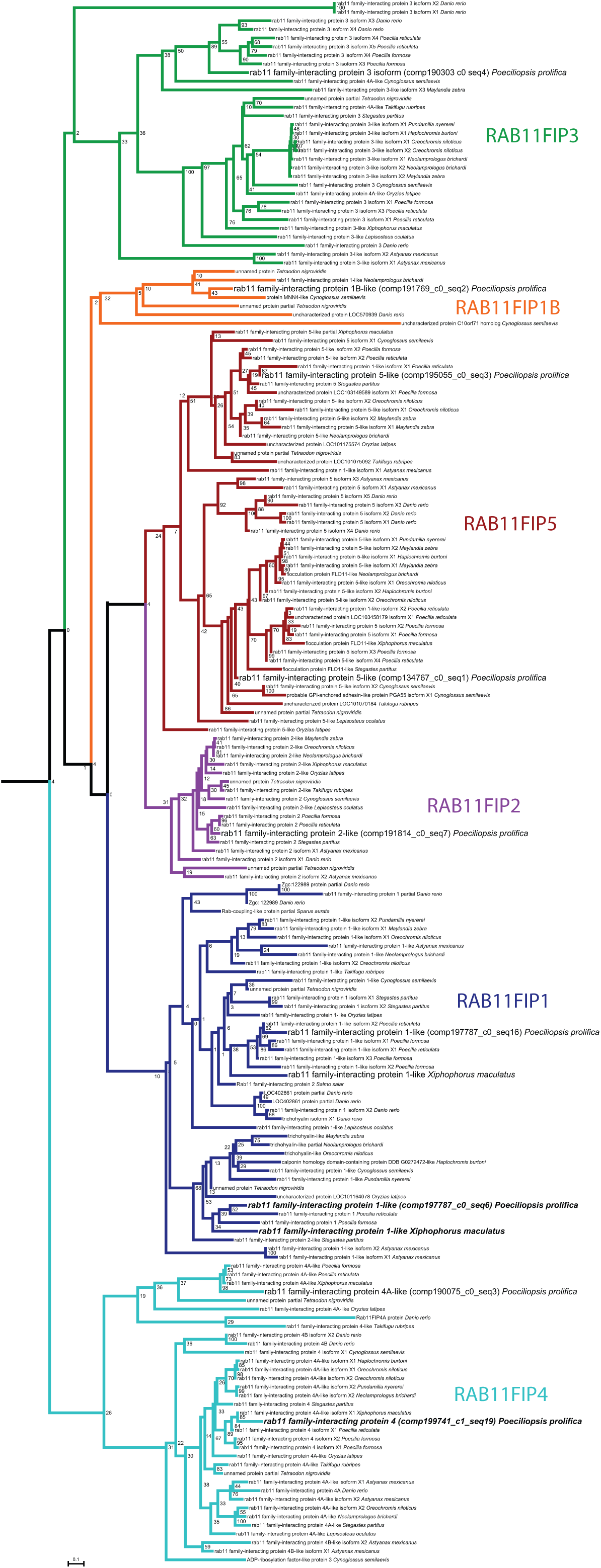
Phylogenetic distance-based gene-family tree for *RAB11 family-interacting proteins* (RAB11FIPs) in fishes. Each color represents different gene family member protein group. RAB11FIPs found in Poeciliopsis prolifica in enlarged fonts. Proteins with sites found to be under positive selection in PAML analysis in bold italics; specifically, RAB11FIP1 in *P. prolifica and X. maculatus* and RAB11FIP4 in *P. prolifica only*.

